# Time-to-target simplifies optimal control of visuomotor feedback responses

**DOI:** 10.1101/582874

**Authors:** Justinas Česonis, David W. Franklin

**Affiliations:** Technical University of Munich, Germany

## Abstract

Visuomotor feedback responses vary in intensity throughout a reach, commonly explained by optimal control. Here we show that the optimal control for a range of movements with the same goal can be simplified to a time-to-target dependent control scheme. We measure participants’ visuomotor responses in five reaching conditions, each with different hand or cursor kinematics. Participants only produced different feedback responses when these kinematic changes resulted in different times-to-target. We complement our experimental data with a range of finite and non-finite horizon optimal feedback control models, finding that only the model with time-to-target as one of the input parameters can successfully replicate the experimental data. Overall, this suggests that time-to-target is a critical control parameter in online feedback control. Moreover, we propose that for a specific task and known dynamics, humans can instantly produce a control signal without any computation allowing rapid response onset and close to optimal control.

## Introduction

From intercepting a basketball pass between opponents to catching a vase accidentally knocked off the shelf – visuomotor feedback responses play a familiar role in human motor behaviour. Previous research has extensively analysed these responses in human reaching movements (*Day and Lyon* (*2000*); *Reichenbach et al.* (*2014*); *de Brouwer et al.* (*2017*, 2018); *Saunders and Knill* (*2003*); *Saunders* (*2004*); *Saunders and Knill* (*2005*); *Sarlegna et al.* (*2003*); *Knill et al.* (*2011*)), and showed an interesting combination of task-dependent variability on the timescale of a single movement (*Dimitriou et al.* (*2013*); *Franklin et al.* (*2014*, 2017)), as well as sub-voluntary feedback onset times (*Prablanc and Martin* (*1992*); *Day and Lyon* (*2000*); *Franklin and Wolpert* (*2008*); *Zhang et al.* (*2018*); *Oostwoud Wijdenes et al.* (*2011*)). These visuomotor feedback responses have been shown to modulate throughout a movement depending on the perturbation onset location (*Dimitriou et al.* (*2013*)). This observation was explained through optimality principles, however such control was modelled only indirectly, by replicating velocity profiles and trajectories of visually perturbed movements (*Liu and Todorov* (*2007*); *Rigoux and Guigon* (*2012*)). In this study we test to what degree optimal feedback control, as opposed to other control methods, can be used to model the visuomotor feedback responses directly.

Optimal control as a theory of human movement has normally been compared against other theories in terms of prediction of kinematics and dynamics (*Todorov and Jordan* (*2002*); *Izawa et al.* (*2008*); *Nagengast et al.* (*2009*); *Yeo et al.* (*2016*); *Guigon et al.* (*2007*, 2008)). Nevertheless, optimal feedback control has been used to motivate extensive studies investigating the control and task-dependent modulation of feedback responses (*Knill et al.* (*2011*); *Pruszynski and Scott* (*2012*); *Nashed et al.* (*2012*, 2014)). The results of these and other studies have highlighted the 2exibility of the modulation of these feedback responses. While a few studies have compared the predictions of the controller feedback gains against the feedback responses in human subjects (*Knill et al.* (*2011*)), such predictions have not been made about the temporal evolution of these feedback responses during reaching. For example, *Dimitriou et al.* (*2013*) show temporal evolution of feedback response intensity throughout a reaching movement, suggesting that this is similar to the feedback gain predictions of *Liu and Todorov* (*2007*). However a direct comparison of these feedback intensities has not been made. Here we directly compare the temporal evolution of visuomotor feedback response intensities in human participants with the prediction of these intensities in an optimal feedback control model.

Visuomotor feedback response intensity over a goal directed reaching movement follows a roughly bell-shaped profile, with peak intensity in the middle and decay towards the beginning and the end of the movement (*Dimitriou et al.* (*2013*)). The results of *Liu and Todorov* (*2007*) suggest that such modulation is a combination of gains related to movement position, velocity and acceleration. However, we do not yet know whether these would be related to the visual or haptic kinematics. In addition, models of ball catching were shown to produce systematic errors in the prediction of the hand kinematics when using only velocity or acceleration based gains (*Dessing et al.* (*2002*)), suggesting an integration of multiple state variables to produce the feedback response. Evidence of such integration then raises two important questions. First, could there be other states than position and its derivatives that also contribute to such control? Second, how can these responses be produced so rapidly, when multiple inputs need to be integrated into one solution?

One method to solve these two problems would be a controller based on time-to-target. Within a state-space system, all state variables are constantly changing with time with a fixed relationship to one another as described by the state transition and control matrices. Such a system can then be re-imagined as a system with time as its input, and these physical states as the hidden states. Such mapping simplifies the multiple input system where the inputs are state variables, to a one-input (time) system. Indeed, the expected time-to-target (or time-to-contact) has been shown to be related to the control in finger pointing (*Oostwoud Wijdenes et al.* (*2011*)) and catching tasks (*Dessing et al.* (*2002*)). Therefore, we test whether a simple relation to the time-to-target can explain the temporal profile of visuomotor feedback responses in humans. To test our hypotheses, we devised an experimental paradigm where we offset the usual bell-shaped velocity profile in the aim to separate the effect of the times-to-target from the effect of kinematics on the visuomotor feedback responses. Finally, we compare these results with a normative optimal feedback control model of visuomotor feedback responses in order to better understand how and whether these responses can be the result of optimality and still maintain rapid onset times.

## Results

### Experimental results

In this study we examine the relation between time-to-target and the visuomotor feedback responses. To do so, we devised an experiment consisting of five different kinematic conditions. The baseline condition required movements with a natural, bell-shaped velocity profile, while the velocity profiles were modified for the four other conditions. In these four conditions we introduced a manipulation between the hand velocity and the cursor velocity in the forward direction, such that the cursor and hand had different velocity profiles, but their positions matched at the start and end of the movement (Figure 1). Two of these four conditions (matched-cursor conditions) required different kinematics of the physical movement to successfully complete the task, but the cursor velocity profiles matched the baseline. This manipulation of hand velocity profiles also resulted in different times-to-target at the same distance in the movement. The two other conditions (matched-hand conditions) required the same hand movement as for the baseline condition, but as a result the cursor moved with different velocity profiles (see Materials and Methods). This manipulation of the cursor velocity profiles separates the relative contributions of physical and visual hand information in regulating the feedback responses. For each condition we measured the visuomotor feedback intensities (mean corrective force applied during 180-230 ms time window after a visual perturbation) at five different locations in the movement (Figure 2A). Overall our paradigm allowed us to modulate the times-to-target across conditions, as well as separate proprioceptive (hand) and visual (cursor) kinematics to examine their individual contribution to visuomotor feedback responses.

**Figure 1.**
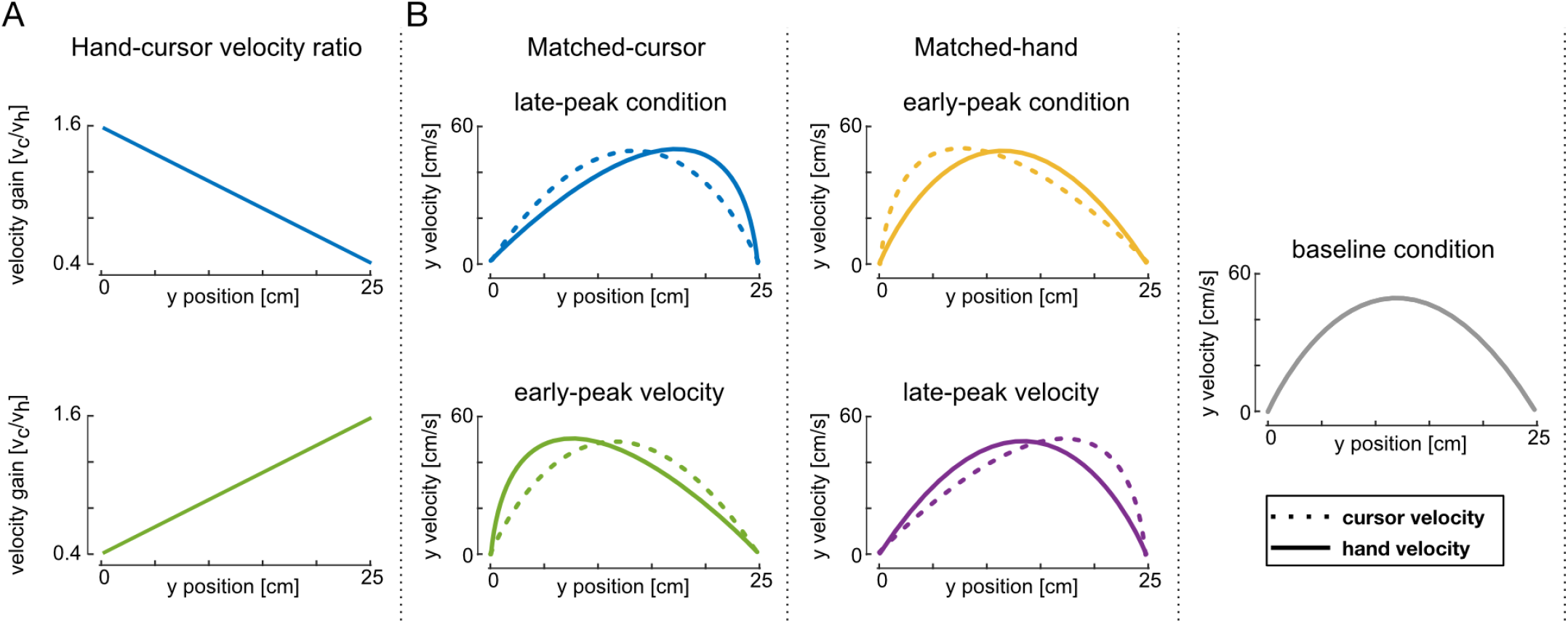
Experimental design. (**A**) Top: hand-cursor velocity scaling for conditions where the cursor position leads the hand position in y axis (matched-cursor late-peak hand velocity condition, blue, and matched-hand early-peak cursor velocity condition, yellow). Bottom: hand-cursor velocity scaling for conditions where the cursor position lags the hand position in y axis (matched-cursor early-peak hand velocity condition, green, and matched-hand late-peak cursor velocity condition, purple). (**B**) Hand and cursor velocity-position profiles required to achieve the ideal movement to the target. Left: matched-cursor velocity conditions; middle: baseline condition, where cursor position and hand position are consistent; right: matched-hand velocity conditions.

**Figure 2.**
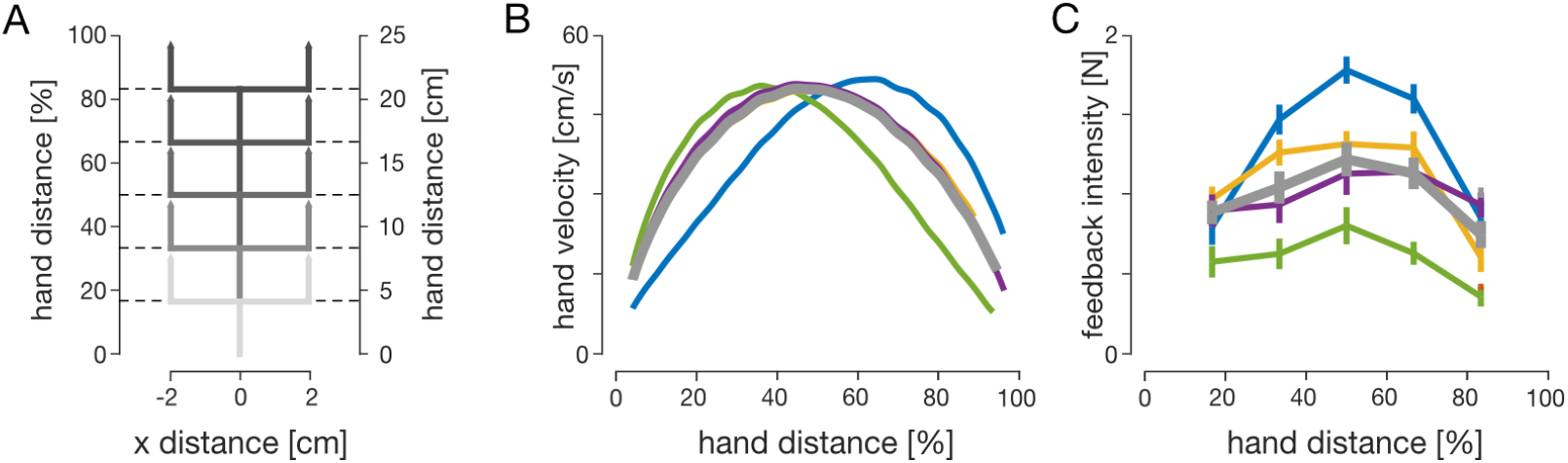
Human visuomotor feedback responses are modulated across the five experimental conditions. (**A**) Lateral perturbations of the cursor were applied in all five conditions. Perturbations were introduced as 2 cm cursor jumps perpendicular to the movement direction. The perturbation onset occurred at one of five equally spaced hand locations. (**B**) Mean velocity profiles of the hand in five experimental conditions: matched-cursor early-peak (green), matched-cursor late-peak (blue), matched-hand early-peak (yellow), matched-hand late-peak (purple) and baseline (grey). Participants successfully modulated forward movement kinematics to meet task demands – velocity profiles are skewed for matched-cursor conditions, and are similar to the baseline for matched-hand conditions. (**C**) Mean visuomotor feedback intensities (mean lateral force from 180-230 ms after perturbation onset) across all participants to cursor perturbations as a function of the hand distance in the movement. Error bars represent 1 SEM. Significant regulation is observed for matched-cursor early-peak and matched-cursor late-peak conditions (blue and green), but no significant regulation is seen for matched-hand conditions (yellow and purple), relative to the baseline.

Different movement conditions exhibited differences in visuomotor feedback intensities (Figure 2 and Figure supplement 1). Two-way repeated-measures ANOVA (both frequentist and Bayesian; Materials and Methods) showed significant main effects for both condition (*F*_4,36_ = 10.807, *p* < 0.001, and *BF*_10_ = 9.136 × 10^12^), and perturbation location (*F*_4,36_ = 33.928, *p <* 0.001, and *BF*_10_ = 6.870 × 10^9^). Post-hoc analysis on movement conditions revealed significant differences between baseline (grey line) and matched-cursor late-peak hand velocity condition (blue line; *t*_9_ = 4.262, *p*_*bonf*_ < 0.001 and *BF*_10_ = 247.868), and between baseline and matched-cursor early-peak hand velocity condition (green line; *t*_9_ = −8.287, *p*_*bonf*_ < 0.001 and *BF*_10_ = 1.425 × 10^8^). However, no significant differences were found between the baseline and the two matched hand velocity conditions (*t*_9_ = 1.342, *p*_*bonf*_ = 1.0 and *BF*_10_ = 0.357 for early-peak cursor velocity, yellow; *t*_9_ = 0.025, *p*_*bonf*_ = 1.0 and *BF*_10_ = 0.154 for late-peak cursor velocity, purple). Our results show that different kinematics of the hand movement have a significant effect on visuomotor feedback response regulation, but that different kinematics of the cursor movement do not.

One possible explanation for differences between the two matched-cursor conditions (blue and green in Figure 2C and Figure supplement 1) and the baseline condition (grey) might arise from a different mapping between cursor and hand velocities (Figure 1A) that had to be learned. Alter-natively, the incongruency between the vision and proprioception might be another explanation. However, the two matched-hand conditions (yellow and purple) had the identical mappings (and incongruencies) as the two matched-cursor conditions (blue and green respectively) and yet no differences were found in these conditions. Instead, the only conditions in which differences in the feedback gains were found, were conditions in which the timing of the peak hand velocity was shifted.

In order to test whether a simple relationship between movement kinematics and visuomotor feedback intensities exists, we mapped visuomotor feedback intensity magnitudes as a linear function of the hand velocity and the cursor velocity. For each experimental condition, we find a different regression slope between the velocity and the feedback intensities regardless of whether this is the cursor or the hand velocity (Figure 3AB). Consistent with our previous results, this difference in slopes is significant for conditions where the hand, but not cursor, movement was different (Figure 3CD). Although feedback intensities increase with increasing velocity in both cursor and hand coordinates, no one coordinate modality could predict the changes in the feedback intensity.

**Figure 3.**
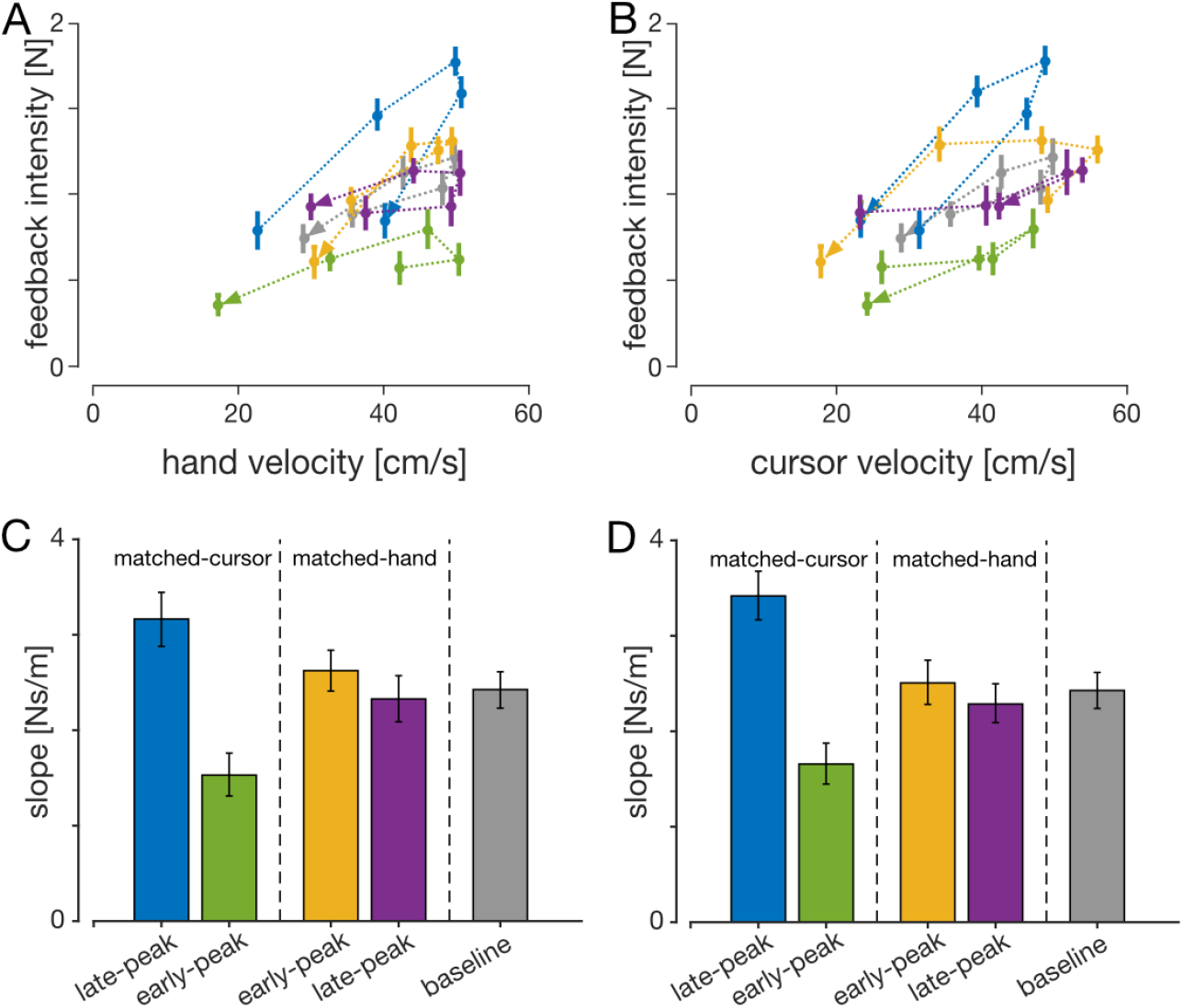
Visuomotor feedback intensities as a function of (**A**) Hand velocity and (**B**) cursor velocity at the time of perturbation for all experimental conditions. Error bars represent 1 SEM, and the arrowheads represent the order of the perturbation locations. (**C**), (**D**) Regression slopes of feedback intensities for each condition as a function of hand and cursor velocities respectively. Error bars represent 95% confidence intervals of the slopes. The slopes for the two matched-cursor conditions were significantly different (based on the confidence intervals) than for the baseline condition.

To successfully complete each trial, participants were required to reach the target. However, the distance to reach the target is affected by the perturbation onset – later perturbation locations lead to larger correction angles (Figure 4A) and thus longer movement distances (Figure 4B). This effect is clearly seen where the extension of movement distance is enhanced for the perturbations closest to the target, with movement distance extended by almost half a centimetre compared to less than one millimetre for the closest perturbations. Any extension of the movement distance requires an appropriate increase in movement duration. Consequently, participants extended their movement time, with longest durations for perturbations close to the target (Figure 5A). This increase in movement duration increases the time-to-target for these late perturbations (Figure 5B), and now allows suZcient time for the controller to issue any corrective commands.

**Figure 4.**
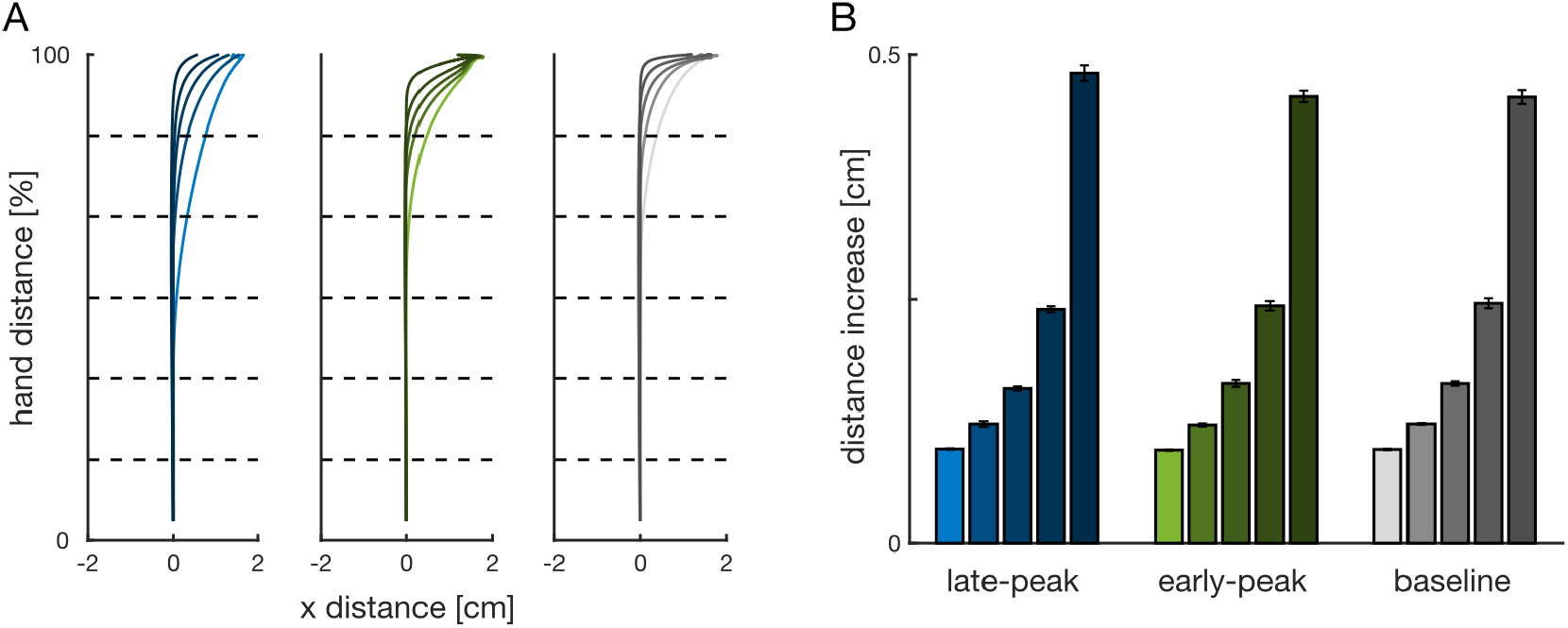
(**A**) Mean hand movement trajectories for matched-cursor late-peak (left), matched-cursor early-peak (middle) and baseline (right) conditions recorded in our participants, with perturbation onset at five locations (colour light to dark: 4.2 cm (16.7%), 8.3 cm (33.3%), 12.5 cm (50%), 16.7 cm (66.7%) and 20.8 cm (83.4%) from the start position; dashed lines). Corrections to rightward perturbations were 2ipped and combined with leftward corrections. (**B**) Distance increase for each perturbation location recorded in our participants. Perturbation locations closest to the target required the largest increases in movement distance. Error bars represent 1SEM.

**Figure 5.**
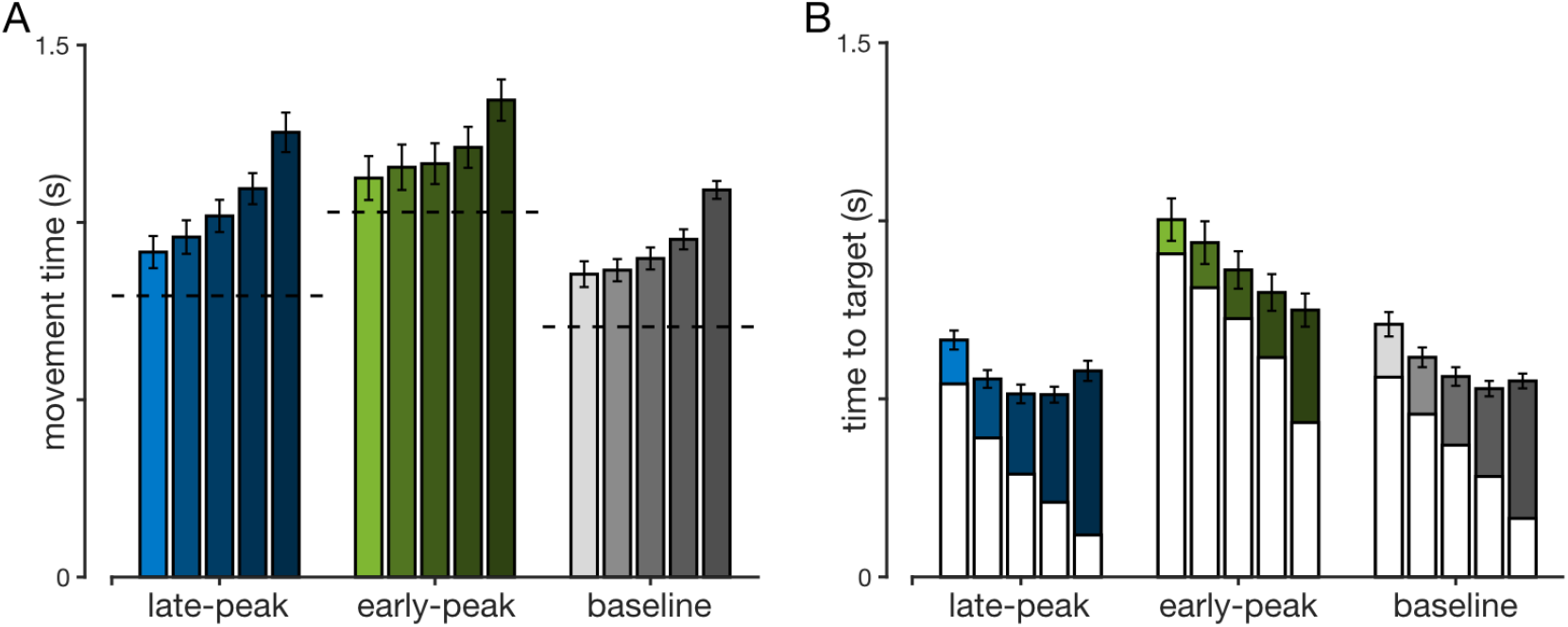
(**A**) Movement durations in maintained perturbation trials recorded by our participants in late-peak, early-peak and baseline conditions. Separate bars within the same colour block represent different perturbation onset locations (left to right: 4.2 cm, 8.3 cm, 12.5 cm, 16.7 cm and 20.8 cm from the start position). Error bars represent 1SEM while the horizontal dashed lines represent movement durations in the same movement condition for non-perturbed movements. (**B**) Full bars represent times-to-target in maintained perturbation trials in our participants for late-peak, early-peak and baseline conditions. White bars represent the time-to-target for a respective non-perturbed movement, at the time when the perturbation would have happened. The coloured part of the bars shows the extension in times-to-target due to the perturbation in a non-constrained movement. Each of the five bars represents a different perturbation onset location, as in (**A**). Error bars represent 1SEM.

### Finite horizon optimal feedback control

As optimal control has been suggested to predict the temporal evolution of feedback intensities (*Dimitriou et al.* (*2013*); *Liu and Todorov* (*2007*)), we built two finite-horizon optimal feedback control (OFC) models: the classical model (*Liu and Todorov* (*2007*)), and a time-to-target model. For the classical model we implemented an OFC (*Todorov* (*2005*)) to simulate movements with different velocity profiles, similar to the experiments performed by our participants. We extended this classical model to the time-to-target model, by increasing the movement duration after each perturbation onset according to experimental results (Figure 5). For both models we only simulated different hand kinematics for computational ease and as our participants showed little effect of cursor kinematics on their feedback intensities.

For both models we controlled the activation cost R to simulate three conditions in which the location of the peak velocity was shifted to match the experimental hand kinematics (Figure 6A). Specifically, we solved for the activation cost R and movement duration N by optimising the log-likelihood of our model’s peak velocity location and magnitude using Bayesian Adaptive Direct Search (BADS, *Acerbi and Ma* (*2017*)). The optimised movement durations (mean ± SEM) were N = 930 ± 0 ms for the baseline condition, N = 1050 ± 10 ms for the late-peak condition and N = 1130 ± 20 ms for the early-peak condition (10 optimisation runs per condition). In comparison, experimental movement durations were N = 932 ± 30 ms for the baseline condition, N = 1048 ± 47 ms for the late-peak condition and 1201 ± 59 ms for the early-peak condition, matching well with the OFC predictions. Overall this shows that specific constraints on the magnitude and location of peak velocity that we imposed on our participants resulted in a modulation of reaching times that matched OFC predictions under the same constraints.

**Figure 6.**
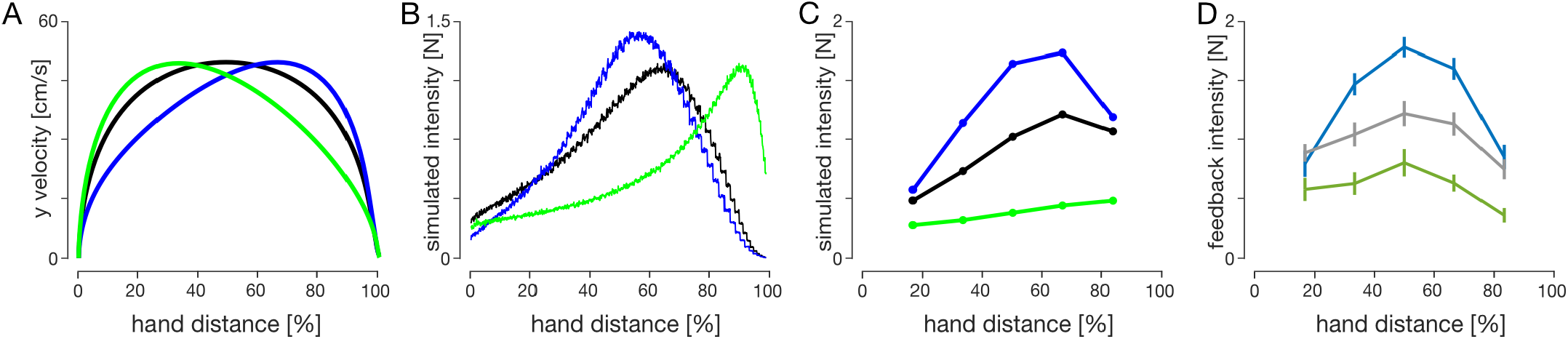
Comparison of feedback intensities between the two OFC models and experimental data. (**A**) Simulated velocity profiles, and (**B**) Simulated feedback intensity profiles of baseline (black), early-peak (green) and late-peak (blue) velocity condition simulations for the classical OFC model. Velocity profiles were obtained by constraining the velocity peak location and magnitude and optimising for movement duration and activation cost function. Simulated feedback intensity profiles were obtained by applying virtual target jumps perpendicular to the movement direction during these movements and calculating the force exerted by the controller in the direction of the target jumps. (**C**) Simulated feedback intensities obtained via the time-to-target OFC model. Pre-perturbation movements were simulated as if no perturbation would occur, in order to keep the controller naive to an upcoming perturbation. At the perturbation onset the remaining movement duration is adjusted to match the mean time-to-target for a similar perturbation onset in human participants (Figure 5B). The velocity profiles for the time-to-target model match the velocity profiles of the classical model, shown in (**A**). (**D**) Visuomotor feedback intensities recorded in human participants.

For the classical model we estimated simulated feedback intensities by shifting the movement target at each timepoint in the movement and measuring the mean magnitude of the simulated force response over a 130-180 ms time window in the direction of this shift. The simulated feedback intensity profiles follow the same general shape as in human participants – intensity increases from the beginning of the movement and then falls off at the end (Figure 6B). However, the overall profile of these simulated feedback intensities is very different for each of the kinematic conditions. For the early-peak velocity condition, the simulated feedback intensity peaks towards the end of the movement (green line), whereas for the late-peak velocity condition the simulated feedback intensity profile peaks early in the movement (blue line). These simulated feedback intensities do not appropriately capture the modulation of visuomotor feedback intensities in our experimental results. Specifically they predict a temporal shift in the peak intensity that is not present in our participants data, and predict similar peak levels of feedback intensities across all three conditions. While the simulated feedback intensities are qualitatively similar to the experimental results within each condition, overall this model cannot appropriately capture the modulation of visuomotor feedback responses across the conditions.

For the time-to-target OFC model, we extended the classical model to account for the different movement durations for each perturbation location (and movement condition) that is seen in the experimental results. After a perturbation, the remaining time-to-target was adjusted to match the experimentally recorded times-to-target for this specific movement, while before the perturbation both the classical model and the time-to-target model were identical. After adjusting for the individual durations of each perturbation condition we are now able to qualitatively replicate the general regulation of feedback intensity profiles for different kinematics using OFC (Figure 6C). In the late-velocity peak condition we predict a general increase in the feedback responses throughout the movement compared to the baseline condition, whereas in the early velocity peak condition we predict a general decrease in these feedback responses compared to the baseline condition. Thus we show that within the OFC the time-to-target is critical for the regulation of feedback responses, and when we take this into account we are able to replicate the feedback intensity modulation of our participants.

While in our experiment, we manipulated the time-to-target through skewing the velocity profiles, time-to-target is naturally modified through changing the peak velocity. Therefore, we can further analyse the effect of the time-to-target by calculating the feedback intensities for movements with different peak velocities (Figure 7A). The simulated feedback intensities vary widely across peak velocities, with a shift of peak feedback intensities towards the earlier locations for faster movements (Figure 7B). However, when these distinct simulated feedback intensity profiles are re-mapped as a function of time-to-target, the simulated feedback intensities follow a consistent, albeit non-monotonic, relationship (Figure 7C). This relationship is also consistent over a range of peak velocities across all three kinematic conditions and is well described by a combination of a square-hyperbolic and logistic function (Figure 7D). The squared-hyperbolic arises from the physics of the system: the lateral force necessary to bring a point mass to a target is proportional to 1/*t*^2^ (Materials and Methods, Equation 9). The logistic function simply provides a good fit to the data. Overall our models show that the feedback intensity profiles under OFC are independent of the peak velocity or movement duration. Instead, our simulations suggest that time-to-target is a key variable in regulating visuomotor feedback responses.

**Figure 7.**
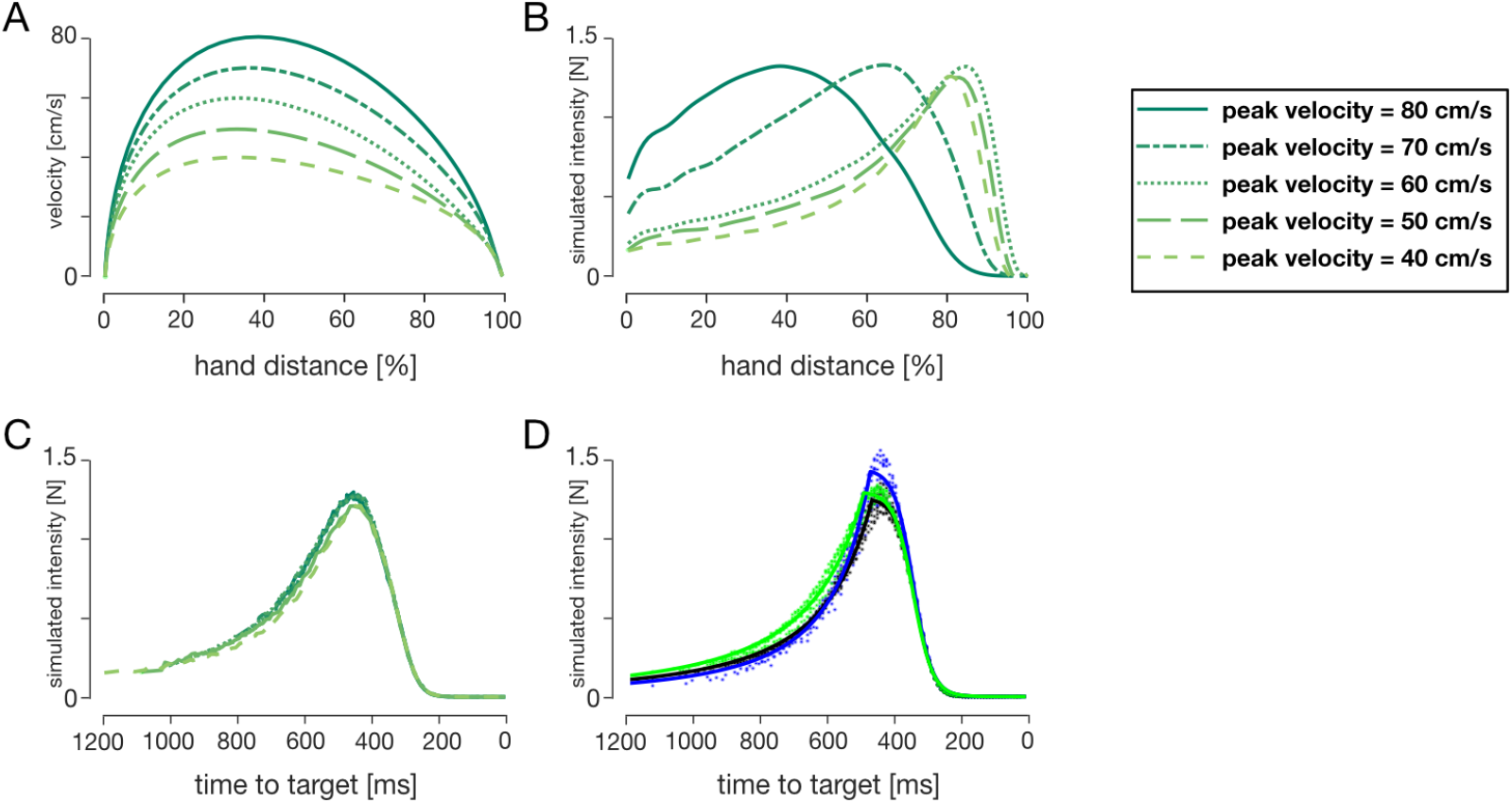
OFC simulations of (**A**) velocity profiles and (**B**) simulated feedback intensity profiles for different desired peak velocities (in order from light to dark line colours: 40 cm/s, 50 cm/s, 60 cm/s, 70 cm/s, 80 cm/s). (**C**) Simulated feedback intensities of (**B**) re-mapped as a function of time-to-target at the time of target perturbation. (**D**) Simulated feedback intensities vs time-to-target for the three kinematic conditions over the five peak velocities simulated by OFC (coloured dots). Solid lines represent the tuning curves (Equation 7) fit to the data. Both the tuning curves and the simulated feedback intensity profiles are similar across a variety of different kinematics when expressed as a function of time-to-target.

It has been shown that the optimal controller gains (*Liu and Todorov* (*2007*)), as well as the visuomotor feedback intensities (*de Brouwer et al.* (*2017*); *Knill et al.* (*2011*)) are influenced by task definition (e.g. instruction to hit the target or stop at the target). Here we simulated the hit, fast hit and stop instructions for our classical model in order to test how it influenced the relation between simulated feedback intensity and time-to-target. Our previous simulations represent the stop instruction. We modified the *ω*_*v*_ and *ω*_*f*_ to simulate the baseline equivalent of hit and fast hit instructions. Specifically, we set *ω*_*v,hit*_ = *ω*_*v*_/4 = 0.05, *ω*_*f,hit*_ = *ω*_*f*_/4 = 0.005 for hit instruction, and *ω*_*v,fasthit*_ = *ω*_*v*_/10 = 0.02, *ω*_*f,fasthit*_ = *ω*_*f*_/10 = 0.002 for fast hit instruction. As changing the terminal costs also results in a change in peak velocity, we further reduced the desired movement times to *N* = 800 ms for the hit instruction and *N* = 750 ms for fast hit instruction, such that all three peak velocities match (Figure 8A). According to our simulations, such modification of task demands produced different simulated feedback intensity profiles (Figure 8B). However, the intensity relationship with time-to-target maintained the same structural profile independent of the task demand (Figure 8C). Specifically, both the squared-hyperbolic and logistic segments of the control are still present, although we observe the shift in the temporal location of the crossover point. While each task requires a different pattern of feedback gains (and will therefore produce different responses), variations of the kinematic requirements within a task do not change these gains and therefore do not require recalculation.

**Figure 8.**
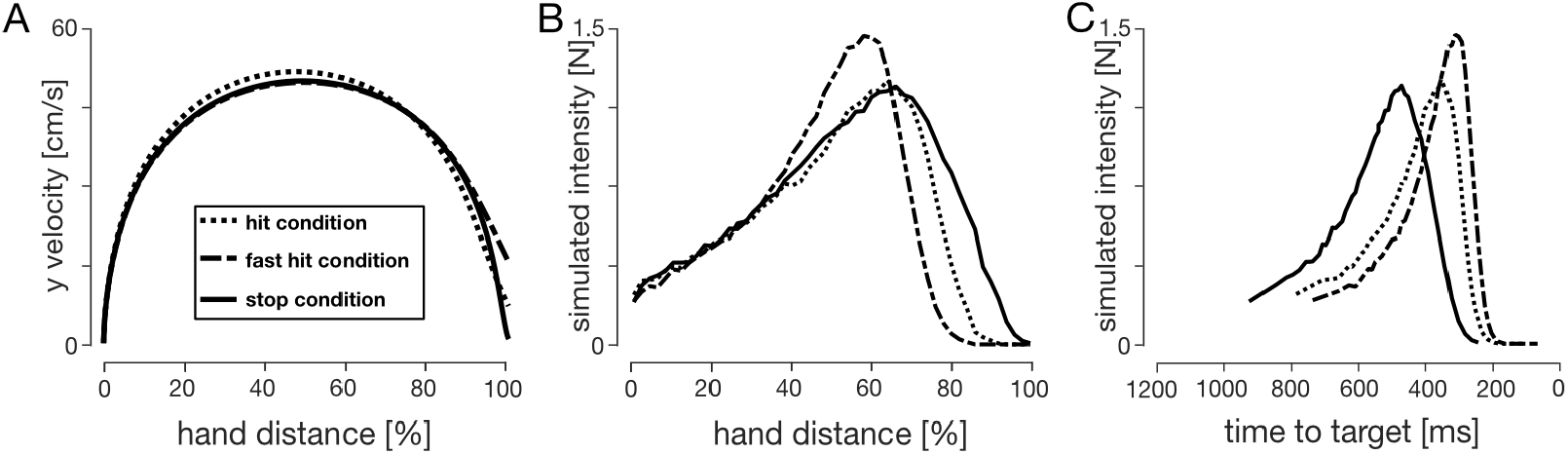
Comparisons between hit and stop instructions. (**A**) Velocity profiles for the stop, hit and fast-hit conditions. (**B**) Simulated feedback intensity profiles as a function of hand position. (**C**) Simulated feedback intensities of (**B**) re-mapped as a function of time-to-target at the time of target perturbation.

### Receding horizon and infinite horizon control

A limitation of the finite-horizon implementation used in classical and time-to-target models is that the variable movement duration (Figure 5) is the model input rather than output. Therefore, in addition to finite-horizon models we also modelled our task in receding and infinite horizon for a single movement condition. Specifically, for the infinite horizon model both state-dependent and regulator costs were kept constant throughout the simulated movement. For the receding horizon model the regulator cost was kept constant, while the state-dependent cost was zero for all but last “foreseeable” state. Such models were expected to simulate the baseline experimental condition, however the resultant velocity profile better resembled the early-peak condition (Figure 9A). As a result, we compared these simulations with both baseline and early-peak velocity condition data and with the time-to-target model simulations (Figure 9B-D).

**Figure 9.**
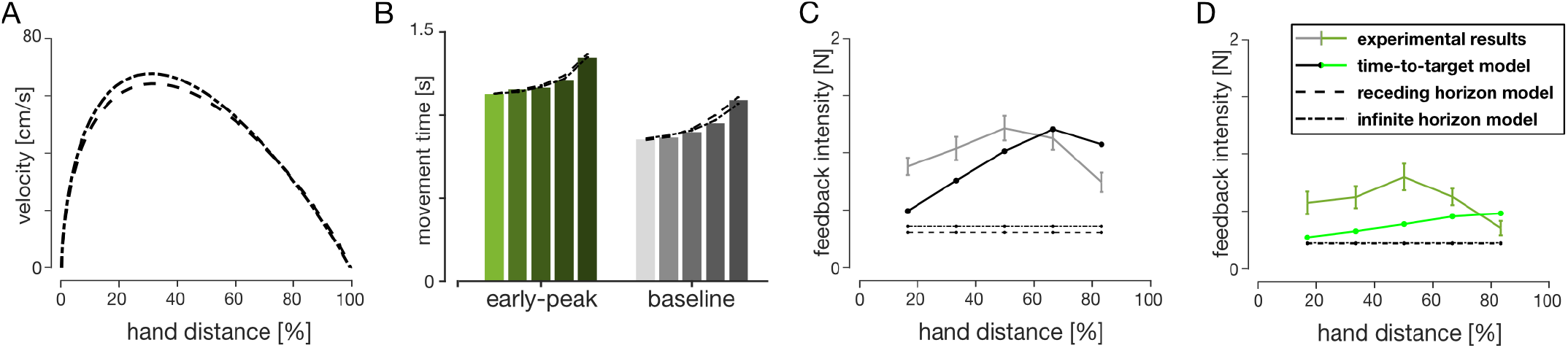
Receding horizon and infinite horizon model simulations. (**A**) Simulated velocity profiles of receding horizon (dashed) and infinite horizon (dot-dashed) models. Both models naturally produce positively skewed velocity profiles, more closely resembling early-peak velocity, rather than the baseline condition. (**B**) Mean experimental movement durations (bar chart) compared to the receding and infinite horizon model predictions. Both models accurately simulate the variations in the reach durations with perturbation location. (**C**) Baseline and (**D**) Early-peak velocity condition simulations for receding horizon, infinite horizon and time-to-target (dot-solid lines) models, compared to the experimental data. Only the time-to-target model predicts different visuomotor feedback response intensities for different perturbation onset locations, while receding and infinite horizon models predict constant intensities. Note that models were not fit to match the intensities, only to qualitatively demonstrate the behaviour.

Both receding horizon and infinite horizon LQG models were able to successfully capture the non-linear change in trial durations for different perturbation onsets (Figure 9B) matching the experimental results. In addition, these models also predicted variable times-to-target for the five perturbation onset locations: (700 ms, 660 ms, 620 ms, 600 ms, 580 ms) for the infinite horizon and (690 ms, 640 ms, 610 ms, 610 ms, 600 ms) for the receding horizon. However, neither model showed variation of the simulated feedback intensities for different perturbation onset locations (Figure 9CD) – a result that was present in the experimental data and captured by our time-to-target model. Instead both models predicted constant feedback intensities for all perturbations locations. Therefore neither the receding nor the infinite horizon models are able to explain our experimental results. While both of the approaches can accurately capture the variability in movement duration, only the time-to-target model well describes the behavioural variation in visuomotor feedback responses.

### Validation of the time-to-target model

Overall our simulations suggest that independent of movement kinematics — different temporal position, velocity, and acceleration profiles — the visuomotor feedback intensities follow the same profile with respect to the time-to-target. We further verified how our time-to-target prediction matches our actual experimental results by plotting participants’ visuomotor feedback intensities against the average time-to-target for the respective perturbation locations and movement conditions (Figure 10A). While we did not specifically fit our time-to-target model to our experimental data, we still see the qualitative similarities between the two. Specifically, the intensities monotonically increase with decreasing time-to-target until the peak (following the squared-hyperbolic function) and then reduce (the logistic function range).

**Figure 10.**
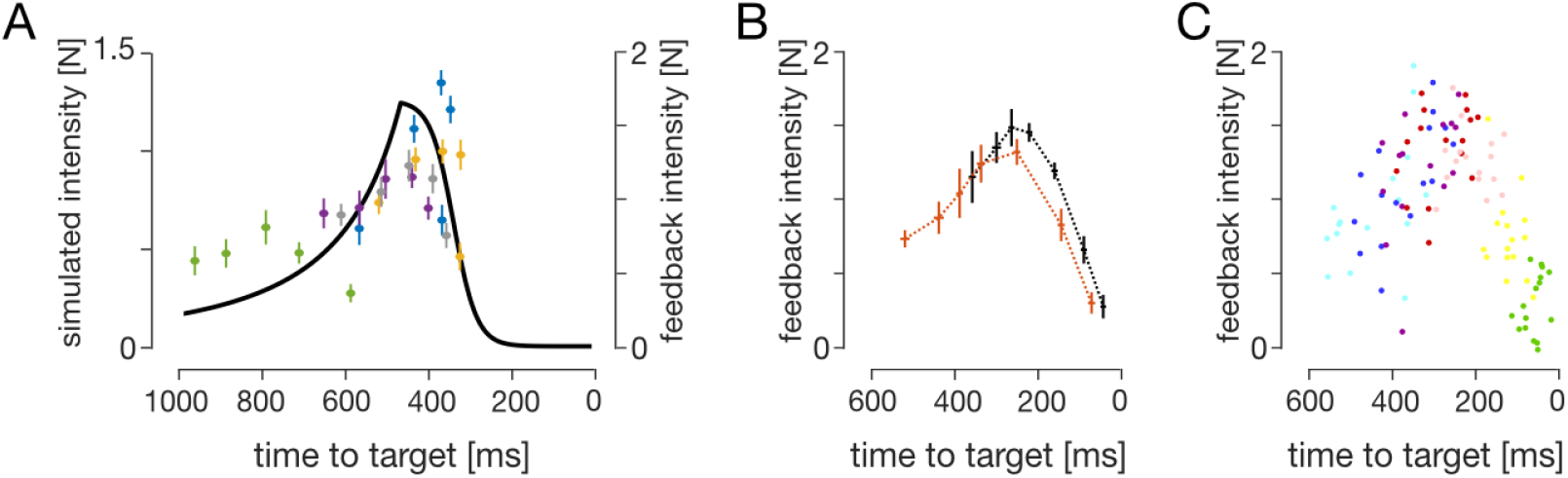
Validation of the time-to-target model. (**A**) Experimental visuomotor feedback intensities for all five experimental conditions (scatter plot) overlaid with the OFC model for the baseline condition, as a function of time-to-target. Error bars represent 1SEM (**B**) Experimental data of the visuomotor feedback intensities of *Dimitriou et al.* (*2013*), mapped against the time-to-target. Black and orange traces represent mean participant data for 17.5 cm and 25 cm movement conditions respectively. (**C**) A scatter plot of individual subjects’ data from (**B**). Different colours represent different perturbation onset distances as in *Dimitriou et al.* (*2013*).

Finally, we also compared the prediction of the time-to-target model to independent results from an external data set (*Dimitriou et al.* (*2013*)). In the article the authors could not rigorously encapsulate both conditions within a simple relationship to movement distance, movement fraction or movement velocity. We plotted visuomotor feedback intensities against time-to-target for two experimental conditions: goal directed reach of 17.5 cm and of 25 cm (Figure 10BC). Two observations can be made from these results. First, the time-to-target model prediction and the experimental data follow the same qualitative features, independent of the target distance (experimental condition). Second, the feedback intensities for both conditions are well explained by a single relationship with time-to-target. All together, both our data and *Dimitriou et al.* (*2013*) data strongly support our time-to-target model.

## Discussion

Here we examined how movement kinematics regulate visuomotor feedback responses. Participants extended their movement duration after perturbations to successfully reach the target. In addition, visuomotor feedback responses were modulated when the hand followed different kinematics, but not when the cursor followed different kinematics. In order to better understand this modulation we built four normative models using OFC: a classical finite-horizon OFC (*Liu and Todorov* (*2007*)), a finite-horizon time-to-target adjusted OFC, a receding-horizon OFC (*Guigon et al.* (*2019*)) and an infinite-horizon OFC (*Qian et al.* (*2013*)). While the classical, receding and infinite horizon models failed to predict the experimental visuomotor feedback response intensities, the time-to-target model qualitatively replicated the visuomotor feedback intensity profile of our participants. Overall, optimal feedback control models suggested that feedback intensities for each perturbation location depended on the time-to-target rather than distance or velocity. Moreover, this explains why any mismatch between visual and haptic kinematics had no effect on the feedback intensities, as these manipulations did not affect the time-to-target. Simulated feedback intensities under all movements followed the same profile with respect to time-to-target, suggesting a critical role in the regulation of visuomotor feedback responses.

Experimentally, our participants exhibited a temporal evolution of visuomotor feedback intensities for each condition, confirming the findings of *Dimitriou et al.* (*2013*). In addition, we also showed the regulation of visuomotor feedback responses across conditions, allowing us to investigate the underlying mechanism of this temporal evolution. Specifically, our experimental results demonstrated strong regulation of visuomotor feedback intensity profiles with different hand kinematics, but not with different cursor kinematics (Figure 2C). Compared to the baseline condition, in the matched-cursor early-peak velocity condition participants produced longer times-to-target at each perturbation location (Figure 5B), resulting in weaker feedback responses based on the relationship between time-to-target and visuomotor feedback intensities (Figure 10A). The opposite is true for the matched-cursor late-peak velocity condition. As the two matched-hand conditions produced similar times-to-target as the baseline due to similar hand kinematics, we did not observe a different regulation in feedback responses. Therefore, the condition-dependent visuomotor feedback response modulation exhibited by our participants meshes nicely with a control policy whereby the time-to-target regulates the feedback responses.

It has long been suggested that we select movements that minimize the noise or endpoint variability (*Harris and Wolpert* (*1998*)). Within the framework of optimal control, this idea has been expanded to the corrective movements – that is, optimality in reaching movements is achieved in part by minimizing the noise during any corrective response (*Todorov and Jordan* (*2002*)). As motor noise scales proportionally to muscle activation (*Jones et al.* (*2002*); *Hamilton et al.* (*2004*)), one way of minimising such noise is reducing the peak levels of muscle activation during the correction. Mathematically, the optimal solution to correct any perturbation approximates a constant activation, resulting in a constant force for the whole duration between perturbation onset and target interception. Such a solution assumes that the brain is capable of estimating the remaining duration of the movement (*Benguigui et al.* (*2003*); *McIntyre et al.* (*2001*); *Zago et al.* (*2004*)) and that the force follows the squared-hyperbolic relationship to this duration (Equation. 9). The parallel can be drawn here between our results and the results of *Oostwoud Wijdenes et al.* (*2011*), where the authors showed a similar temporal evolution of peak acceleration against the time-to-target in a single forward velocity condition. Our results further show that time-to-target strongly modulates visuomotor feedback responses across a range of different kinematics, consistent with the idea that human participants aim to behave optimally. More specifically, we suggest that, among different optimality variables, the temporal evolution of visuomotor feedback response intensities serves to reduce effects of system noise.

Finite-horizon OFC predicts a time beyond which feedback responses are suppressed. Beyond this critical time, a logistic function well describes the relation between time-to-target and feedback responses, with response intensities reducing as the time-to-target decreases. The controller gains at this stage are the most sensitive to acceleration, suggesting a more “behavioural” outcome – the controller is trying to stop, rather than correct errors. The neural recordings in rhesus macaque monkeys’ supplementary motor area and M1 (*Russo et al.* (*2019*)) show that SMA can signal movement termination as far as 500 ms before the end of the movement. This further suggests that there may be multiple stages within a movement, where our control system might “care” more about error correction in one or movement termination in another. On the other hand, the suppression of responses close to the target leads to undershooting the target. Our participants, however, had to bring the cursor to the target in order to advance to the next trial. As a result, they extended the movement durations post-perturbation to return to the squared-hyperbolic range of control. The control performance of such behaviour is well accounted for by our time-to-target model. Moreover, our time-to-target model also well explained the modulation of visuomotor feedback intensities from an external data set (*Dimitriou et al.* (*2013*)). However, an important distinction from our study is that in *Dimitriou et al.* (*2013*) the suppression of feedback responses towards the end of movements would not interfere with reaching the target as perturbation trials were always in a mechanical channel so that no corrections were required. As a result, the times-to-target were shorter and the data clearly exhibits both logistic and squared-hyperbolic segments of the control.

A limitation of our time-to-target model is that it takes time-to-target as an input in order to generate feedback intensity predictions, rather than obtain the time-to-target as a model output. As a result, our time-to-target model does not describe exactly how the change in movement geometry after the perturbation influences this time-to-target, which in turn regulates the visuomotor feedback responses. On the other hand, both receding and infinite horizon models did predict the movement duration change after perturbations very well, but could not at all describe the changes in visuomotor response intensity. However, utility of movement has recently been used within optimal control to characterise reaching movements (*Rigoux and Guigon* (*2012*); *Shadmehr et al.* (*2016*)) in which optimal movement time falls out automatically from a trade-off between reward and effort. With respect to our models, this adds additional complexities to capturing the different movement conditions. Future approaches could attempt to model these results within the utility of movement framework.

In addition, our time-to-target model does not directly show the causality of the time-to-target as a control variable for the visuomotor feedback intensities. Particularly, the time-to-target relation to feedback intensity could be a by-product of a more sophisticated control scheme. Additional arguments for the time-to-target control scheme could be two-fold. First, there is evidence that humans are well capable of estimating the time-to-target of a moving stimulus, even if it is accelerating (*Benguigui et al.* (*2003*); *McIntyre et al.* (*2001*); *Zago et al.* (*2004*)), indicating that time-to-target is at least an available input for such a controller. Second, while we have tested finite-horizon OFC and two other (receding and infinite horizon) OFCs, only the finite horizon controllers had any effect on the variation of simulated feedback intensities. Importantly, neither the receding nor infinite horizon models use time-to-target as an input to the controller. We posit that this time-to-target control input is the one key difference between the finite and non-finite models and is therefore the simplest explanation for our results.

Rapid feedback responses scale with the temporal urgency to correct for mechanical perturbations (*Crevecoeur et al.* (*2013*)). Here we have shown that visuomotor feedback responses also follow a similar regulation, suggesting that these two systems share the same underlying control policy. Our work further extends this finding of Crevecoeur by not just showing that temporal urgency affects feedback responses, but explaining the manner in which these responses are regulated with respect to urgency. That is, here we have shown that for visual perturbations the feedback intensities scale with a squared-hyperbolic of the time-to-target, which is a direct measure of urgency. Moreover, the feedback intensities were rapidly adjusted due to the change in urgency as the task changed. Specifically, when the cursor jumps close to the target, the expected time-to-target is prolonged, and therefore the optimal visuomotor feedback response needs to be adjusted appropriately to this increase in time. Our results show that participants produce a visuomotor response consistent with the actual, post-perturbation, time-to-target, as opposed to the expected time-to-target prior to the perturbation. Therefore, our results not only suggest that similar computations might occur for both stretch and visuomotor feedback response regulation, but also that this regulation originates from task-related optimal feedback control.

Our work has shown that simulated feedback intensities from OFC exhibit the same underlying pattern as a function of time-to-target over a wide range of movement kinematics, matching well the feedback intensities of our human participants (Figure 6). As expected, changes in the task goals (e.g. hit versus stop) changed the relation between feedback responses and time-to-target. However, the qualitative features – the squared-hyperbolic and logistic function – remained consistent across these tasks. These results suggest that, for a specific task and known dynamics, we do not need to recalculate the feedback gains prior to each movement, but instead can access the appropriate pattern as a function of the estimated time-to-target in each movement. Therefore gain computation in reaching movements may not be a computationally expensive process, but instead could be part of an evolutionary control strategy that allows for rapid estimation of the appropriate feedback gains. Moreover, the fact that both stretch reflex and visuomotor feedback systems exhibit similar control policies despite different sensory inputs, perhaps only sharing the final output pathway, suggests that this simple feedback pathway may be an evolutionary old system. Indeed, several studies have suggested that visuomotor feedback is controlled via a pathway through the colliculus (*Reynolds and Day* (*2012*); *Gu et al.* (*2018*); *Corneil et al.* (*2004*)). Such a system would then only need to be adapted as the dynamics or overall task goals change, allowing for fine tuning of the feedback gains according to changes in the environment (*Franklin et al.* (*2017*)).

Our results have shown the connection between the visuomotor feedback response regulation and the time left to complete the movement. Specifically, in our human participants we recorded the increase in the time-to-target after the perturbation onset, which consequently increased the movement durations (Figure 5). This increase was also longer for later perturbations, consistent with previous studies (*Liu and Todorov* (*2007*)). According to our normative time-to-target OFC model, the time-to-target alone is enough to successfully regulate visuomotor feedback responses as observed in humans. This result was independent of the kinematics of the movement or the onset times of the perturbations. This suggests that there is no recalculation of a control scheme for the rest of the movement after the perturbation, but rather a shift to a different state within the same control scheme. Such findings are consistent with the idea that visuomotor feedback gains are pre-computed before the movement, allowing for faster than voluntary reaction times (*Franklin* (*2016*)). Moreover, through our results, we gain a deeper insight into how optimal feedback control governs these feedback gains – through a straightforward relationship to the estimated time-to-target, based on physics.

## Materials and Methods

### Participants

Eleven right-handed (*Oldfield* (*1971*)) human participants (5 females; 27.3 ± 4.5 years of age) with no known neurological diseases took part in the experiment. All participants provided written informed consent before participating. All participants except one were naïve to the purpose of the study. Each participant took part in five separate experimental sessions, each of which took approximately 3 hours. One participant was removed from analysis as their kinematic profiles under the five experimental sessions overlapped. The study was approved by the Ethics Committee of the Medical Faculty of the Technical University of Munich.

### Experimental setup

Participants performed forward reaching movements to a target while grasping the handle of a robotic manipulandum with their right hand. Participants were seated in an adjustable chair and restrained using a four-point harness. The right arm of participants was supported on an air sled while grasping the handle of a planar robotic interface (vBOT, *Howard et al.* (*2009*)). A six-axis force transducer (ATI Nano 25; ATI Industrial Automation) measured the end-point forces applied by the participant on the handle. Position and force data were sampled at 1kHz. Visual feedback was provided in the plane of the hand via a computer monitor and a mirror system, such that this system prevented direct visual feedback of the hand and arm. The exact onset time of any visual stimulus presented to the participant was determined from the graphics card refresh signal.

Participants initiated each trial by moving the cursor (yellow circle of 1.0 cm diameter) into the start position (grey circle of 1.6 cm diameter) located approximately 25 cm in front of the participant, centred with their body. This start position turned from grey to white once the cursor was within the start position. Once the hand was within the start position for a random delay drawn from a truncated exponential distribution (1.0-2.0 s, mean 1.43 s), a go cue (short beep) was provided signalling participants to initiate a straight reaching movement to the target (red circle of 1.2 cm diameter, located 25.0 cm directly in front of the start position). If participants failed to initiate the movement within 1000 ms the trial was aborted and restarted. Once the cursor was within 0.6 cm of the centre of the target, participants were notified by the target changing colour to white. The movement was considered complete when the participants maintained the cursor within this 0.6 cm region for 600 ms. If participants did not complete the movement within 4 seconds from first arriving at the start position (e.g. by undershooting or overshooting the target), the movement timed-out and had to be repeated. After each trial, the participant’s hand was passively returned by the robot to the start position while visual feedback regarding the success of the previous trial was provided (Figure 11). Movements were self-paced, and short breaks were enforced after every 100 trials.

**Figure 11.**
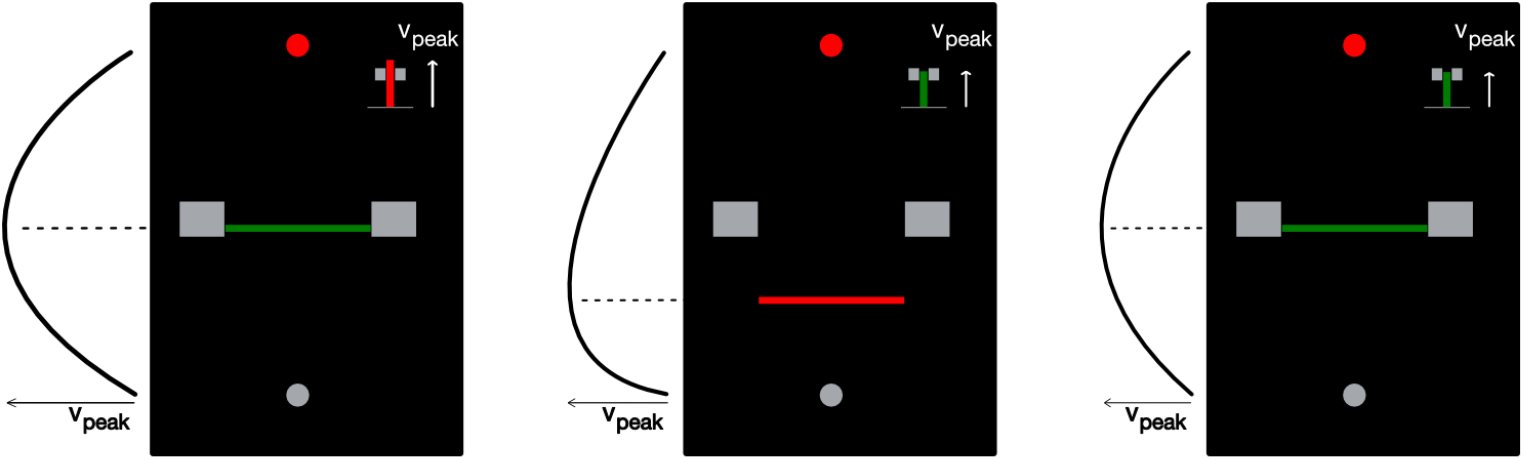
Examples of feedback presented to the participants. Feedback regarding the peak velocity and the timing of the peak velocity was provided after each trial. Large grey blocks indicate the velocity peak location target, while the bar chart at the top-right corner indicates peak y-velocity magnitude. Feedback was provided on the modality (cursor or hand) that matched the baseline. Left: velocity peak location is within the target, but the movement was too fast (unsuccessful trial); middle: velocity peak location is too early, but the movement speed is within the target (unsuccessful trial); right: successful trial.

### Experimental paradigm

Participants performed the experiment under five different conditions, each performed in a separate session. In the baseline condition the cursor matched the forward movement of the hand, with a peak velocity in the middle of the movement. In the other four conditions, the cursor location was scaled relative to the hand location in the forward direction, such that the cursor and the hand location matched only at the start and end of the movements (Figure 1). In two of the conditions (matched-hand velocity), the hand velocity matched the baseline condition throughout the movement (with the peak in the middle of the movement) but the cursor velocity peaked either earlier (33% of movement distance) or later (66% of movement distance). In the other two conditions (matched-cursor velocity), the cursor velocity was matched to the baseline condition throughout the movement (with the peak in the middle of the movement) but the hand velocity peaked either earlier (33% of movement distance) or later (66% of movement distance). The difference between the cursor velocity and the hand velocity was produced through a linear scaling of the cursor velocity as a function of the forward position (Figure 1A). Specifically, for the two conditions where the position of the peak cursor velocity is earlier than the position of the peak hand velocity (Figure 1 top), this scaling was implemented as:

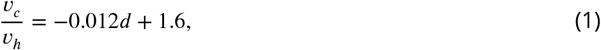

where *v*_*c*_ and *v*_*h*_ are cursor and hand velocities respectively, and *d* is the distance along the movement direction in %. The cursor velocity was therefore manipulated by a linear scaling function such that its velocity is 160% of the hand velocity at the beginning of the movement, linearly decreasing to 40% at the target location (Figure 1 top). For the two conditions where the position of the peak cursor velocity is later than the position of the peak hand velocity (Figure 1 bottom), this scaling was implemented as:

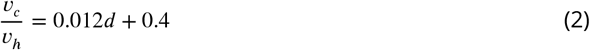

such that the velocity gain function linearly increased from 40% hand velocity at the start of the movement to 160% at the end of the movement (Figure 1, bottom). Desired velocity profiles of both the hand and the cursor are shown in Figure 1B for each condition.

### Feedback regarding movement kinematics

In all conditions, one of the velocity modalities (cursor or hand) was required to be similar to the baseline velocity profile. Feedback was always provided about this specific velocity modality. Ideal trials were defined as trials in which this peak velocity was between 42 cm/s and 58 cm/s with the peak location between 45% and 55% of the movement distance with no target overshoot. After each trial, visual feedback about the peak velocity and the location at which this peak occurred was provided to the participants graphically (Fig 11). The peak velocity was indicated on the right hand side of the screen with the length of a bar and the velocity target. This bar changed colour from red to green if the velocity was within the ideal range. The location of the peak velocity was indicated as a horizontal line between home and target positions at the exact location it was achieved, along with the ideal range. This line was green when the location of the peak velocity was within the ideal range, and red otherwise. Overshooting the target was defined as the position of the cursor exceeding the centre of the target in the y-coordinate by more than 0.9 cm. If participants reached the target while overshooting during the movement, a message indicating the overshot was shown, no points were scored and an error tone was played in order to discourage further overshots.

### Probe trials

During each session, probe trials were used to measure the visuomotor feedback intensity – the average strength of corrective motor response to a change in the visual feedback of hand position. To elicit these feedback responses (further visuomotor feedback responses), visual perturbations were initiated laterally (±2.0 cm) at five different hand distances (4.2, 8.3, 12.5, 16.7, and 20.8 cm) from the start (Figure 2A). In addition a zero amplitude perturbation (cursor matched to the lateral position of the hand) was included, resulting in eleven different probe trials. On these trials the visual perturbations lasted 250 ms, after which the cursor was returned to the lateral location of the hand. The lateral hand position was constrained in a simulated mechanical channel throughout the movement, thereby requiring no correction to reach the target. The simulated mechanical channel was implemented with a stiffness of 4000 N/m and damping of 2 Ns/m acting perpendicularly to the line connecting the start position and the target (*Scheidt et al.* (*2000*); *Milner and Franklin* (*2005*)), allowing measurement of any lateral forces in response to a visual perturbation.

In previous experiments, feedback response intensity gradually decreased during the course of the experiment (*Franklin and Wolpert* (*2008*); *Franklin et al.* (*2012*)). However, it has been shown that including perturbation trials where the perturbations were maintained until the end of the movement, and where participants had to actively correct for the perturbation to reach the target, prevents this decrease in the feedback intensity (*Franklin et al.* (*2016*)). Therefore half of the trials contained the same range of perturbations as the probe trials but where these perturbations were maintained throughout the rest of the trial and participants had to correct for this perturbation. These maintained perturbations have now been used in several studies *Franklin et al.* (*2016*, 2017); *de Brouwer et al.* (*2017*).

### Session design

Prior to each session, participants performed 100 to 300 training trials in order to learn the specific velocity profiles of the reaching movements. All training trials contained no visual perturbations and were performed in the null force field. The training trials were stopped early once participants achieved an accuracy of 75% over the last 20 trials, and were not used for the analysis.

Each session consisted of 40 blocks, where each block consisted of 22 trials performed in a randomized order. Eleven of these 22 trials were probe trials (5 perturbation locations × 2 perturbation directions + zero perturbation condition) performed in the mechanical channel. The other eleven trials consisted of the same perturbations but maintained throughout the trial and performed in the null field. Therefore in each of the five sessions participants performed a total 880 trials (440 probe trials). The order of the five different conditions (sessions) was pseudo-randomized and counterbalanced across participants.

### Data analysis

Data was analyzed in MATLAB R2017b and JASP 0.8.2. Force and kinematic time series were low-pass filtered with a tenth-order zero-phase-lag Butterworth filter (40 Hz cutoff). The cursor velocity was calculated by multiplying the hand velocity by the appropriate scaling function. The visuomotor feedback response was measured for each perturbation location as the difference between the force responses to the leftward and rightward perturbations within a block. To measure the visuomotor feedback response intensity (mean force, produced as a response to a fixed-size visual perturbation) this response was averaged over a time window of 180-230 ms, a commonly used time interval for the involuntary visuomotor feedback response (*Franklin and Wolpert* (*2008*); *Dimitriou et al.* (*2013*); *Franklin et al.* (*2012*, 2016)). In order to compare any differences across the conditions a two-way repeated-measures ANOVA was performed with main effects of condition (5 levels) and perturbation location (5 levels). As a secondary method to frequentist analysis we also used the Bayesian factor analysis (*Adrian E. Raftery and Robert E. Kass* (*1995*)) to verify our statistical results. Bayesian factor analysis is a method that in addition to the conventional hypothesis testing (evaluating evidence in favour of the alternative hypothesis) allows us to evaluate evidence in favour of the null hypothesis, therefore distinguishing between the rejection of the alternative hypothesis and not enough evidence to accept the alternative hypothesis.

Although we used the time window of 180-230 ms to estimate visuomotor feedback intensity, we also verified whether the onset of the visuomotor feedback response in our data is consistent with previously reported values. To estimate this onset time, we first estimated individual onset times for each participant at each perturbation location and movement condition. To do so, we used the Receiver Operator Characteristic (ROC) to estimate where the force reaction to leftwards cursor perturbations deviated from the reaction to rightwards cursor perturbations (*Pruszynski et al.* (*2008*)). For each type of trials we built the ROC curve for the two signals at 1 ms intervals, starting from 50 ms before the perturbation, and calculated the area under this curve (aROC) for each of these points until the aROC exceeded 0.75 for ten consecutive milliseconds. In order to find where the force traces start deviating from each other we then fit a function of the form *max*(0.5, *k* × (*t* − *τ*) to the aROC curve. The time point where the linear component of this function first overtakes the constant component was taken as the threshold value. Overall, the mean onset times across all conditions and perturbation locations were 138 ± 7 ms (mean + SD), with onset times consistent among movement conditions (*F*_4,36_ = 1.410, *p* = 0.25, and *BF*_10_ = 0.105), perturbation locations (*F*_4,36_ = 1.582, *p* = 0.20, *BF*_10_ = 0.252), and their interactions (*F*_16,144_ = 1.350, *p* = 0.176, and *BF*_10_ = 0.005)

### Modelling

#### Optimal feedback control

In addition to our linear models we implemented two different Optimal Feedback Control (OFC) models: the classical model (*Liu and Todorov* (*2007*)) and the time-to-target model. In both models we modelled the hand as a point mass of *m* = 1.1 kg and the intrinsic muscle damping as a viscosity *b* = 7 Ns/m. This point mass was controlled in a horizontal plane by two orthogonal force actuators to simulate muscles. These actuators were controlled by the control signal *u*_*t*_ via a first order low-pass filter with a time constant *τ* = 0.05 s. The state-space representation of the dynamic system used to simulate the reaching movements can be expressed as

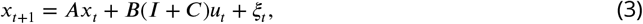

where *A* is a state transition matrix, *B* is a control matrix, and C is a 2 × 2 matrix whose each element is a zero-mean normal distribution representing control-dependent noise. Variables *x*_*t*_ and *u*_*t*_ are state and control at time t respectively. State *x*_*t*_ exists in the Cartesian plane and consists of position **p** (2 dimensions), velocity **v** (2), force **f** (2) and target position **p**^*^ (2). The presence of these four states within the state vector means that the information about all of these states is eventually used for the control. For our simulation purposes we treat the control-independent noise *ξ*_*t*_ as zero.

The state of the plant is not directly observable, but has to be estimated from noisy sensory information. We model the observer as

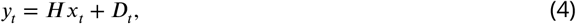

where *H* = diag[1,1,1,1,1,1,0,0] is the observation matrix, and *D*_*t*_ is a diagonal matrix of zero-mean normal distributions representing state-independent observation noise. Therefore, our observer can infer the state information of position, velocity and applied force of the plant, consistent with human participants.

The simulated movements were guided by the LQG controller with a state-dependent cost Q, an activation cost R, a reaching time N, and a time step t = 0.01 s. However, due to the presence of the control-dependent noise, the estimation and control processes are not anymore separable as in the classic LQG theory. In order to obtain optimal control and Kalman gain matrices we utilised the algorithm proposed by *Todorov and Li* (*2005*) where control and Kalman gain matrices are iteratively updated until convergence.

For both the classical and time-to-target models we simulated three different movement kinematics representing three different conditions in our experiment – the baseline and the two matched-cursor conditions. The state-dependent cost Q was identical for all three kinematics:

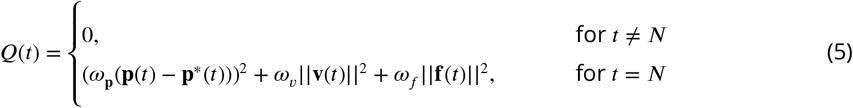

where *ω*_**p**_ = [0.5,1], *ω*_*v*_ = 0.02, and *ω*_*f*_ = 2. The activation cost R(*t*) = 0.00001 was constant throughout the movement for the baseline condition, but was modulated for the two matched-cursor conditions by multiplying it elementwise by a scaling function:

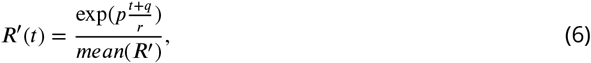

where *p*, *q* and *r* are constants.

Thus, each movement condition only differed from the other two by the profile of this activation cost R, but not by its magnitude. These modified activation costs shift the timing of the peak velocity towards either the beginning or the end of the movement by penalising higher activations at either the end or beginning of the movements respectively. The mean activation cost is kept constant across the conditions resulting in each condition being equally “effortful”. All other simulation parameters were kept constant across the three conditions.

Although LQG is a fixed time horizon problem, we did not pre-define the movement duration N. Instead, we obtained the N, and constants *p*, *q* and *r* using Bayesian Adaptive Direct Search (BADS, *Acerbi and Ma* (*2017*)) to maximise the log-likelihood of the desired peak velocity location and magnitude. We did not fit any other parameters beyond this point. Rather, we analysed our models’ qualitative behaviour compared to human participant data.

The classical and the time-to-target models only differed in the way the perturbations were handled. For the classical model, we simulated perturbation trials at every time step *t*_*p*_ by shifting the target x-coordinate by 2 cm at the time *t*_*p*_ + 120 ms. This 120 ms delay was used in order to mimic the visuomotor delay in human participants, and was taken from *Liu and Todorov* (*2007*). We then averaged the force response of the controller over the time window [*t*_*p*_ + 130, *t*_*p*_ + 180] as an estimate of the simulated feedback responses, equivalent of visuomotor feedback responses in our participants. This means that our simulated feedback responses arise due to separate contributions from the controller position, velocity and acceleration gains. For perturbations occurring at times where the movement is over before the end of this time window, the intensity of this simulated feedback response is set to zero.

For the time-to-target model we introduced an extension in the time-to-target after the onset of any perturbation similar to that observed in our participants. Simulated feedback intensities were modelled at five locations, matching the perturbation locations in our experiment to obtain the appropriate increase in time-to-target after each perturbation. In order to simulate the response to perturbations we first extracted the perturbation onset times from movement kinematics by performing an unperturbed movement and recording the timepoint *t*_*p*_ at which this movement passed the perturbation onset location. We then simulated the post-perturbation portion of the movement as a new LQG movement with an initial state matching the state at *t*_*p*_ + 120 ms of the unperturbed movement, and movement duration matching the time-to-target recorded in our participants for the particular perturbation. Together this keeps our simulated reaches “naive” to the perturbation prior to its onset and allows the time-to-target of the simulated reaches to match the respective time-to-target of our human participants. Finally, we calculated the simulated feedback intensities as described previously, using a time window [10 ms, 60 ms] of the post-perturbation movement. As in the previous simulations, these simulated feedback responses arise due to separate contributions from the controller position, velocity and acceleration gains.

### Time-to-target tuning function

In order to understand the mechanisms that might underlie the consistent relationship between the simulated feedback intensities and the time-to-target, we fit a mathematical expression to the simulated feedback intensities. We modelled the relationship as the minimum of a squared-hyperbolic function and a logistic function:

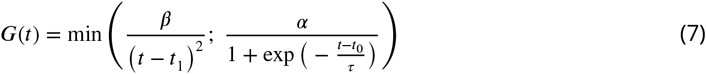

and used BADS to fit this function to our time-to-target–simulated feedback intensity data (Fig. 7C) by optimising the log-likelihood of this fit.

While the logistic function was chosen simply as it provided a good fit to the data, the squared-hyperbolic arises from the physics of the system. Specifically, from the kinematic equations of motion for a point mass (*m*) travelling a distance (*d*) under the influence of force *F*, the distance can be expressed as:

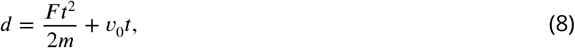

where *v*_0_ = 0 is the lateral velocity at the start of perturbation correction. Rearranging gives:

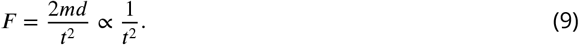

Hence the lateral force necessary to bring a point mass to the target is proportional to 1/*t*^2^.

### Receding horizon OFC

In addition to our finite horizon control we also implemented a receding horizon controller (*Guigon et al.* (*2019*)). Irrespectively of the current state of the movement *X*_*t*_, the receding horizon controller is defined to aim to arrive at the target at time *t* + *T*_*h*_. In essence, such controller is therefore not different from the finite horizon controller in its implementation for a single state of the movement. We implemented the receding horizon controller by iterating a finite horizon controller described previously, but with the *T*_*h*_ = 500 ms, and Q and R costs scaled from the finite horizon model to fit the movement duration. For each iteration we recorded the next movement state (10 ms away from the initial state), and used that as the initial state for the next iteration. This process was repeated until the cursor was within the distance of 0.4 cm from the target position, and remained there without overshooting for 600 ms.

Simulating differently skewed velocity profiles within the framework of receding-horizon control is non-trivial. As a result, we chose to only model one, the baseline, experimental condition, where the activation cost R is constant within the movement. Therefore we chose the costs

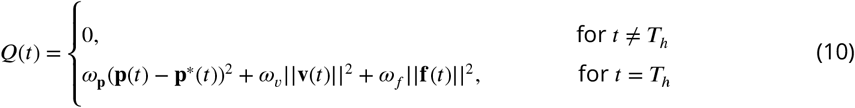

where *ω*_**p**_ = [5,5], *ω*_*v*_ = 0.05, and *ω*_*f*_ = 5. and the activation cost R = 0.000003. The values were selected so that the movement durations, produced by the receding-horizon model would match the experimental durations for the baseline condition (Figure 5A). However, the resultant velocity profiles of this model more closely resembled those of the early-peak velocity condition, than those of the baseline. To account for any effects of the velocity profile we also fit the costs so the model prediction of movement durations matched the durations of the early-peak velocity condition. For this simulation we selected *ω*_**p**_ = [0.7,0.7], *ω*_*v*_ = 0.007, and *ω*_*f*_ = 0.7, while the activation cost remained unchanged.

In this model we introduced the simulated perturbation by shifting the target position by 2 cm at 120 ms after the y-coordinate of the movement passed the perturbation onset location. We only simulated the perturbations matching our experimental conditions–lateral 2 cm cursor jumps, with the onset at five evenly distributed forward distances. We calculated simulated feedback intensities the same way as for the classical and time-to-target models.

### Infinite horizon OFC

We implemented the infinite horizon OFC to control our simulated hand based on the previous work of *Qian et al.* (*2013*). Specifically, we calculated the control gain matrix L, and Kalman gain matrix K to control the same system as in the previous models. We chose the state-dependent costs *ω*_**p**_ = [1,1], *ω*_*v*_ = 0.02, and *ω*_*f*_ = 0 for the baseline condition simulation, and *ω*_**p**_ = [0.35,0.35], *ω*_*v*_ = 0.007, and *ω*_*f*_ = 0 for the early-peak condition simulation. For both conditions the activation cost R=0.002 was kept the same. The protocol of simulating the mean trajectories, feedback responses and their intensities was otherwise identical to the receding horizon simulations.

## Acknowledgements

We thank Matthew Millard, Michael Dimitriou, Sae Franklin and Raz Leib for their comments on an earlier version of this manuscript.

## Notes

#### Summary of Updates

Revisions to the manuscript including the motivation, removing linear modelling section, adding infinite and receding horizon optimal control models and new simulation tasks.

