## Supplemental Figure 1 for "Time-to-target simplifies optimal control of visuomotor feedback responses"

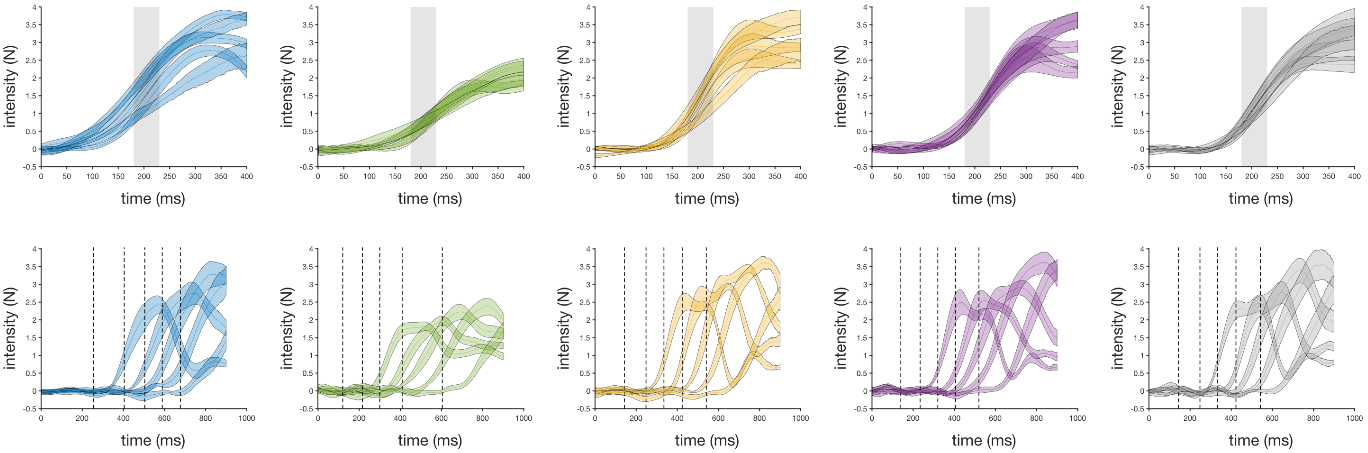


**Figure supplement 1.** Visuomotor feedback response profiles of the five experimental conditions. **Top**: visuomotor feedback responses, aligned to the perturbation onset time. The force response exhibits an onset time of around 130 ms after the perturbation. Visuomotor intensity is calculated by averaging the visuomotor response over a time window of 180-230 ms (shaded area). The colours of the response traces represent different conditions as described in Figure 1B. Shaded areas represent 1SEM. **Bottom**: visuomotor feedback responses, aligned to the movement onset time. Dashed lines represent the perturbation onset time for each different perturbation onset location. The colours of the response traces represent different conditions as described in Figure 1B. Shaded areas represent 1SEM.
